# Properties of individual hippocampal synapses influencing NMDA-receptor activation by spontaneous neurotransmission

**DOI:** 10.1101/590141

**Authors:** Sarah R. Metzbower, Yuyoung Joo, David R. Benavides, Thomas A. Blanpied

## Abstract

NMDA receptor (NMDAR) activation is critical for maintenance and modification of synapse strength. Specifically, NMDAR activation by spontaneous glutamate release has been shown to mediate forms of synaptic plasticity as well as synaptic development. Interestingly, there is evidence that within individual synapses each release mode may be segregated such that postsynaptically there are distinct pools of responsive receptors. In order to examine potential regulators of NMDAR activation due to spontaneous glutamate release in cultured rat hippocampal neurons, we utilized GCaMP6f imaging at single synapses in concert with confocal and super-resolution imaging. Using these single spine approaches, we found that Ca^2+^ entry activated by spontaneous release tends to be carried by GluN2B-NMDARs. Additionally, the amount of NMDAR activation varies greatly both between synapses and within synapses, and is unrelated to spine and synapse size, but does correlate loosely with synapse distance from the soma. Despite the critical role of spontaneous activation of NMDARs in maintaining synaptic function, their activation seems to be controlled factors other than synapse size or synapse distance from the soma. It is most likely that NMDAR activation by spontaneous release influenced variability in subsynaptic receptor position, release site position, vesicle content, and channel properties. Therefore, spontaneous activation of NMDARs appears to be regulated distinctly from other receptor types, notably AMPARs, within individual synapses.

**Significance Statement:** Understanding the underlying synaptic mechanisms for learning and memory is critically important to the field of neuroscience and for human health. A key neurotransmitter receptor type involved in learning is the NMDA receptor, and exploration of its regulation is vital. In this study, we optimized optical tools to allow detailed characterization of NMDA receptor activity at single synapses, along with analysis of structural features of the imaged synapses. The amount of receptor activation is independent of the size of the synapse, but weakly dependent on synapse position within the dendritic tree. Notably, we found that NMDA receptors activated following spontaneous neurotransmitter release tend be GluN2B-containing receptors. Thus, the unique mechanisms that regulate the number and positioning of these receptors within synapses will have important consequences for control of synaptic development and signaling.

## Introduction

NMDA-type glutamate receptor (NMDAR) activation is critical for learning and memory. This is due to the unique ability of NMDARs to allow influx of Ca^2+^ into dendritic spines following glutamate binding and sufficient membrane depolarization (MacDermott et al., 1986; Ascher and Nowak, 1988). It is well established that NMDAR activation by action potential (AP)-dependent neurotransmitter release initiates complex signaling within the postsynaptic density (PSD), which underlies many forms for synaptic plasticity and is essential for normal synaptic function (Kennedy, 2000; Hardingham et al., 2001; Hardingham and Bading, 2003; Higley and Sabatini, 2012; Huganir and Nicoll, 2013). However, NMDAR activation by spontaneous glutamate release, while less studied, also plays a central role in synaptic function. NMDAR-mEPSCs in hippocampus are quite small, ranging from 3 to 20 pA (Prybylowski and Wenthold, 2004; Watt et al., 2004; Prybylowski et al., 2005), and they show a high degree of variability in amplitude (Bekkers and Stevens, 1989). Nevertheless, blockade of NMDAR-mEPSCs induces local protein synthesis-dependent synaptic plasticity (Sutton et al., 2004; Frank et al., 2006; Sutton et al., 2007; Aoto et al., 2008), and NMDAR-mEPSCs mediate synaptic development at distal sites (Andreae and Burrone, 2015). It has been hypothesized that the amount of Ca^2+^ influx through NMDARs within a synapse may lead to different downstream effects (Cummings et al., 1996) and therefore is a critical factor in determining how individual synapses function. Thus, understanding how NMDAR activation due to spontaneous release events is regulated at individual synapses is essential for our understanding of how synapses are maintained and modulated.

Though spontaneous transmission clearly utilizes mechanisms that for the most part are in common with action potential-evoked transmission, some evidence has accrued to suggest that it involves unique elements both pre and postsynaptically. Application of open channel blockers during evoked release led to an elimination of responses due to evoked but not spontaneous release and vice versa for NMDARs, as well as AMPA receptors. This suggests that receptors activated by spontaneous release may form a synaptic pool separate from receptors activated by evoked release within a single synapse (Atasoy et al., 2008; Sara et al., 2011). Additionally, there is some evidence that glutamate vesicles that fuse spontaneously are drawn from a specific subset of the total vesicle pool (Sara et al., 2005; Fredj and Burrone, 2009; Ramirez and Kavalali, 2011; Andreae et al., 2012), though this is still a subject of some debate (Prange and Murphy, 1999; Groemer and Klingauf, 2007; Hua et al., 2010; Wilhelm et al., 2010). However, that spontaneous activation drives signaling pathways through NMDAR activation at individual synapses that are unique from those induced by evoked release (Sutton et al., 2007; Autry et al., 2011), supports the notion that a portion of synaptic NMDARs are specifically activated by spontaneously released glutamate. Taken together this highlights the need to understand the factors at individual synapses that control the type and numbers of NMDARs activated by spontaneous release.

One such parameter could be NMDAR subunit composition, which is a major factor that influences both receptor biophysical properties and downstream signaling pathways (Paoletti et al., 2013; Shipton and Paulsen, 2014b). NMDARs are typically assembled from two GluN1 subunits and two GluN2 subunits that either are two GluN2A (GluN2A-NMDARs), two GluN2B (GluN2B-NMDARs), or triheteromers with one of each type of GluN2 subunit (GluN2A/B-NMDARs) (Rauner and Köhr, 2011; Tovar et al., 2013; Hansen et al., 2014). These receptor types have biophysical differences in glutamate affinity, channel open time, and decay kinetics that lead to differences in Ca^2+^ influx magnitude and kinetics (Santucci and Raghavachari, 2008). In addition, they engage in different protein-protein interactions within the PSD to gate unique downstream signaling pathways (Paoletti et al., 2013; Shipton and Paulsen, 2014b). Thus, which NMDAR subunits contribute to NMDAR activation by spontaneous release may underlie differences in receptor activation and subsequent functional effects.

For AMPA receptors (AMPARs), there is a strong relationship between the amount of receptor activation and the size of the spine (Matsuzaki et al., 2001). This suggests that synapse structure may be an additional critical factor in modulating NMDAR activation by spontaneous release. However, neither the number of NMDARs present (Kharazia et al., 1999; Takumi et al., 1999; Shinohara et al., 2008; Chen et al., 2015), nor the amount of NMDAR activation by glutamate uncaging (Sobczyk et al., 2005) or evoked release (Nimchinsky et al., 2004), was correlated with synapse size. Nevertheless, because NMDAR activation due to spontaneous release may diverge from the total synaptic NMDAR pool, we asked whether spontaneous NMDAR activation is sensitive to synaptic spine or synapse size.

Here we combined single-synapse Ca^2+^ imaging with confocal and super-resolution microscopy to explore the relationship between NMDAR activation due to spontaneous single vesicle exocytosis and morphological features of individual synapses. We find that NMDAR activation by spontaneous release is mediated primarily through GluN2B-NMDARs. The amount of Ca^2+^ influx was not correlated with spine or synapse size, and only weakly correlated with synapse position on the neuron. However, the magnitude of activation was highly variable at individual synapses. Thus, the high degree of variability of spontaneous NMDAR activation is most likely dominated by stochastic channel fluctuations and by nanoscale intrinsic properties of the synapse including receptor position, release site position, and the glutamate concentration profile following each release event.

## Materials and Methods

### Cell culture and GCaMP6f expression

Dissociated hippocampal cultures were prepared from E18 rats of both sex and plated on glass coverslips coated with poly-A-lysine. A subset of cells was infected with pAAV.CAG.GCaMP6f.WPRE.SV40 (Penn Vector Core) at DIV0 and plated along with uninfected cells so that on each coverslip there is a mix of infected and uninfected cells. This allowed us to vary the ratio of infected cells to uninfected cells plated based on the requirement of the experiment. For most experiments 10-15K infected cells were plated with 40K uninfected cells to have a total density ∼50K/well. For correlative Ca^2+^ imaging and super-resolution experiments we used 1K infected cells plated with 30K uninfected cells per well. Transfections were performed using lipofectamine transfection at DIV18 with either GCaMP6f or tdTomato, and constructs were allowed to express for three days, followed by imaging on DIV21. All animal procedures were performed in accordance with the [Author University] animal care committee’s regulations.

### Ca^2+^ imaging

Ca^2+^ imaging was performed on three very similar spinning disk confocal systems. The first was a CSU-22 confocal (Yokagawa) with a Zyla 4.2 sCMOS camera (Andor) mounted on the side port of an Olympus IX-81 inverted microscope, using a 60×/1.42 oil-immersion objective. The second system utilized the same camera, microscope, and objective as above but the CSU-22 confocal was replaced with a Dragonfly multimodal system (Andor). All time-lapses were acquired at 20 Hz controlled by IQ3 (Andor). Following time-lapse acquisition, a z-stack of the field was acquired using 50-200 ms frames and 0.5 µm steps between planes. Coverslips were imaged in ACSF containing 0 mM Mg^2+^, 139 mM NaCl, 2.5 mM KCl, 1.5 mM CaCl_2_, 10 mM glucose, and 10 mM HEPES, pH adjusted to 7.4 with NaOH, 1 μM TTX (Enzo), 10 μM DNQX (Sigma), 20 μM ryanodine (Tocris), 1 μM thapsigargin (Sigma) and 5 μM nifedipine (Sigma). All experiments were performed using an objective heater to maintain bath temperature near 37°C. In order to maintain the plane of focus, autofocus was performed every minute using an Olympus ZDC2. For experiments with a treatment (adding blockers, ifenprodil, raising Ca^2+^, and raising Mg^2+^), baseline imaging of 4-6 minutes was followed by application of either drugs or vehicle (ACSF or DMSO, depending on drug) and then by an additional 4-6 minutes of imaging. The third spinning disk confocal system was a Nikon W1 equipped with a 40x/1.3 oil immersion objective and Hamamatsu Flash4.2 camera, mounted on a Ti2 microscope. This was used for imaging the larger dendritic trees in Figure 5, and images were acquired using otherwise the same parameters and conditions as above. For experiments in Extended Data Figure 2.1, experiments using transfected cells, as well as D-serine experiments were performed on a Nikon TI2 equipped for widefield fluorescence imaging using a 40x/1.3 oil immersion objective and a Zyla 4.2 sCMOS camera. Autofocus was maintained continuously, and acquisition parameters were otherwise the same as above.

### Ca^2+^ Imaging Analysis

In order to analyze the Ca^2+^ imaging data, averages of the first 50-100 frames were generated either in Matlab or MetaMorph. On each averaged image, a circular region of interest (ROI) was drawn around every single spine that was in focus, distinct from the dendrite, and unobstructed, regardless of activity level, as well as a background ROI. Mean intensity within each region was measured for every frame using custom Matlab scripts. Background subtraction was done by subtracting the average intensity of the background ROI from the average intensity of each spine ROI per frame. In order to calculate ΔF/F, F_baseline_ was determined for each spine ROI every minute by averaging fluorescence intensity every 10 frames, and within every minute interval of imaging finding the lowest positive value. Then for each frame (F_frame_ − F_baseline_)/ F_baseline_ was calculated. In order to detect and measure peaks, the ΔF/F values were then fed into Clampex (Axon Instruments) where mSCaTs were detected using a template search that identified peaks based on an average shape profile. For measurements normalized to baseline, only spines that had at least three mSCaTs at baseline were included, typically ∼23-55% of all spines imaged. While for measurements that are not normalized, all spines are included for frequency data and spines with at least one mSCaT were used for amplitude data.

### Spine and cell morphological analysis

Spine area was measured using maximum projections of the post-Ca^2+^ imaging GCaMP6f z-stacks in MetaMorph. Within each spine ROI, an intensity-based threshold was used to calculate area. In order to measure synapse distance from the soma and branch depth, GCaMP6f z-stacks were imported into Imaris (Bitplane) for semi-automatic dendrite and spine detection. Spines identified in Imaris were matched to Ca^2+^ imaged spines using custom Matlab scripts.

### Immunocytochemistry

For immunocytochemistry, cultured hippocampal neurons were fixed in 4% paraformaldehyde (PFA) and 4% sucrose directly following Ca^2+^ imaging for 20 minutes at room temperature (RT). Following fixation, coverslips were washed with Phosphate buffered saline (PBS) + Glycine and permeabilized in PBS+ 0.3% Triton X-100 for 20 minutes at RT. Next, cells were incubated in blocking buffer containing 10% Donkey Serum (DS) (Sigma) and 0.2% Triton X-100 for 1 hour at room temperature. For labeling of Shank, cells were incubated in 1:200 anti-Shank primary (NeuroMab; RRID:10672418), 5% DS and 0.1% Triton X-100 for 3 hours at RT. Followed by incubation with 1:200 Alexa-647 conjugated goat anti-mouse secondary antibody (Jackson) for 1 hour at RT. Finally, cells were post-fixed in 4% PFA and 4% sucrose for 10 minutes.

### Super-resolution imaging

dSTORM imaging was performed on the Olympus IX-81 inverted microscope, using a 60x/1.42 oil-immersion objective along with Dragonfly multimodal system (Andor) and an iXon+ 897 EM- CCD camera (Andor). Cells that had undergone Ca^2+^ imaging were first located by eye in PBS, which was then replaced with STORM imaging buffer containing 50 mM Tris, 10mM NaCl, 10% glucose, 0.5 mg/ml glucose oxidase (Sigma), 40 µg/ml catalase (Sigma), and 0.1 M cysteamine (Sigma) before imaging. Acquisition was performed at 20 Hz, for a total of 70,000 frames while autofocusing every 1000 frames. Imaging was carried out in widefield mode using a power density of 4 was used to concentrate the excitation beam. This allowed for more efficient bleaching of the Alexa-647 molecules.

### Super-resolution imaging analysis

All dSTORM analysis was done using custom Matlab (Mathworks) scripts that fit peaks with an elliptical 2D Gaussian function to a 5 by 5 pixel array. The fitted peaks were used to determine x and y coordinates of the molecules. Molecules with a localization precision less than 20 were used for analysis. PSD detection was performed using custom Matlab scripts. Briefly, following localization detection and drift correction, the image was re-rendered with 14.75 nm pixels, and clusters of localizations exceeding the density cutoff of 1 localization per 217.6 nm^2^ were identified. Clusters with areas less than 0.02 μm^2^, the bottom of the range reported for synapses imaged with super-resolution (MacGillavry et al., 2013), were rejected. Spine Ca^2+^ data was matched to individual PSDs by overlaying super resolved PSDs on the GCaMP6f z-stack and manually matching spines between the post-STORM z-stack and the Ca^2+^ imaging time-lapse.

### Immunoblotting

Hippocampal neuron cultures were prepared as described, infected on DIV0 with AAV vectors, and plated at density of ∼100,000 cells per well in 12-well plate. Neurons were matured as described and on DIV12 or DIV19, neuronal cell homogenates were prepared in preheated buffer containing 1% SDS, 50 mM NaF, 1 mM sodium orthovanadate, and Phosphatase Inhibitor Cocktail I (Sigma) and Phosphatase Inhibitor Cocktail III (Sigma) supplemented with protease inhibitors (Roche cOmplete, EDTA-free). Homogenates were collected into microcentrifuge tubes, sonicated using manual ultrasonication probe, and heated at 65C for 10 min. Protein concentration was determined by BCA assay (Pierce) and equal amounts of total protein (10 micrograms) from each sample were resolved by SDS-PAGE and transferred to PVDF membranes for detection by near-infrared fluorescence. Total protein stain was performed on all membranes (REVERT, LI-COR Biosciences). Immunoblot images were obtained on Odyssey Imaging System and quantitated using Image Studio 4.0 software (LI-COR Biosciences). NMDAR subunit protein levels were normalized against total protein signal for each lane. Quantitative plots were constructed using normalized mean values from 6 samples collected from each condition on each maturation day from a single culture. Data represent mean values +/- standard error of the mean (SEM). Statistical analyses were conducted on quantitated values using GraphPad software.

### Antibodies

Membranes were blotted with primary antibodies including rabbit polyclonal anti-GluN1 C-terminus (1:2000, Sigma-Aldrich Cat# G8913, RRID:AB_259978), rabbit polyclonal anti-GluN2A N-terminus (1:500, JH6097 gift from R. Huganir), mouse monoclonal anti-GluN2B C-terminus (1:2000, Millipore Cat# 05-920, RRID:AB_417391), and mouse monoclonal anti-α Tubulin (1:2000, Sigma-Aldrich Cat# T6074, RRID:AB_477582). Near-infrared conjugated secondary antibodies were used to detect signal for GluN1 or GluN2A: Donkey anti-Rabbit 680RD (1:10,000, LI-COR Biosciences Cat# 926-68073, RRID:AB_10954442), GluN2B: Donkey anti-Mouse 800CW (1:10,000, LI-COR Biosciences Cat# 926-32212, RRID:AB_621847), and α- Tubulin: Goat anti-Mouse 680RD (1:10,000 LI-COR Biosciences Cat# 926-68070, RRID:AB_10956588).

### Experimental Design and Statistical Analysis

Unless otherwise stated, the number of spines was greater than 200; however, treatment with ifenprodil or Mg^2+^ reduced event frequency so that there were fewer spines to use for mSCaT amplitude measurements post-treatment and unless otherwise stated these came from at least 3 separate culture preparations. Statistical analysis was performed using GraphPad Prism. Data are shown as mean ± SEM. Krusal-Wallis followed by Dunn’s test for multiple comparisons were used for experiments that had more than two groups, otherwise Student’s t-test was used to compare means at p<0.05. For experiments comparing the effect of a treatment between groups, post-treatment parameters were normalized to each spine’s own baseline in order to assess the impact of the treatment. For correlations, Pearson’s correlation coefficients were used to assess the strength of the relationship. When noted, data was binned into 8 bins so that pattern in the data could be observed more clearly, however no statistics were performed on binned data. Additionally, for data represented by violin plots, outliers were removed using the ROUT method of identifying outliers. However, all statistics were performed on the raw data before outlier removal.

## Results

### Measuring NMDAR-mediated Ca^2+^ transients with GCaMP6f at individual synapses

To assess NMDAR-activation by spontaneous neurotransmitter release, we sparsely infected dissociated rat hippocampal neurons at the time of plating and imaged neurons at 19- 22 DIV (unless otherwise noted) in ACSF containing 0 mM Mg^2+^, and TTX (1 μM) to block APs. Clear <u>m</u>iniature <u>s</u>pontaneous <u>Ca</u>^2+^ transients (mSCaTs) were detected in individual spines (Fig. 1A) that were well isolated within spines and did not correspond to increases in Ca^2+^ in dendrites (Fig. 1B). Though Ca^2+^ influx through NMDARs can lead to Ca^2+^ release from intracellular stores (Reese and Kavalali, 2015), we found that mSCaT amplitude in spines were unaffected by ryanodine and thapsigargin to prevent Ca^2+^-induced Ca^2+^ release (CICR) from internal stores (vehicle: 0.8164 ± 0.032 ΔF/F, n=277 spines/7 neurons; blockers: 0.9206 ± 0.055 ΔF/F, mean ± SEM, n=239/6; p= 0.095, unpaired t-test) (Fig. 1E, F). Despite this, in order to ensure isolation of the NMDAR component of spine Ca^2+^ changes, all experiments were done in the presence of DNQX to block AMPARs, nifedipine to block L-type Ca^2+^ channels, and previously mentioned CICR blockers. Application of the NMDAR antagonist APV nearly eliminated all events (post-treatment event frequency normalized to baseline: vehicle: 0.9369 ± 0.03228, n=495/10, APV: 0.05541 ± 0.008763, n=197/5; p<0.0001, unpaired t-test) (Fig. 1G, H), confirming that the observed Ca^2+^ transients were NMDAR-mediated.

**Figure 1:**
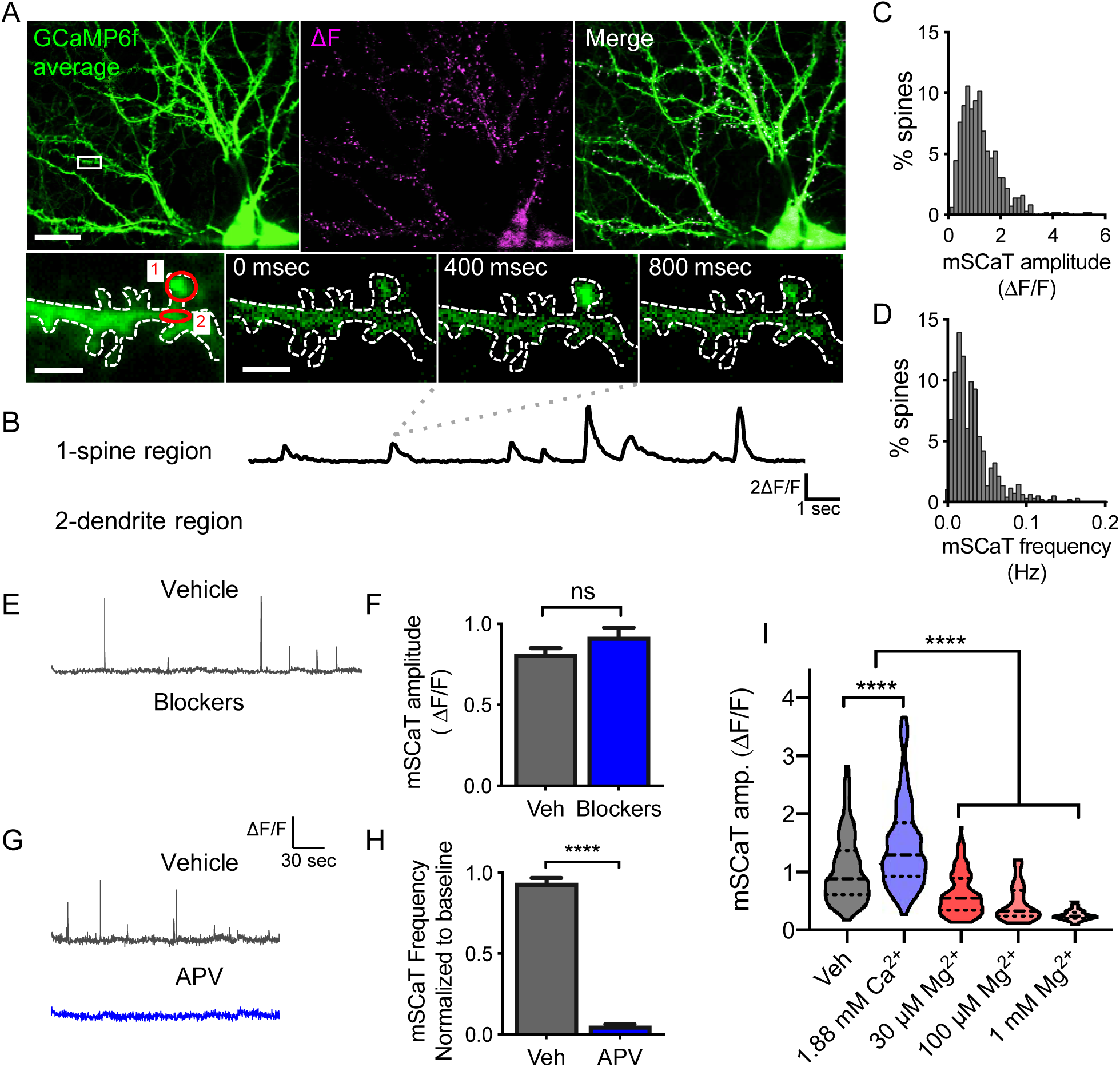
mSCaTs measured by GCaMP6f imaging reflect NMDAR activation at individual synapses following spontaneous single vesicle release. A. Cultured hippocampal neuron infected with AAV-GCaMP6f. Left panel: GCaMP6f average (green), middle panel: ΔF calculated by subtracting GCaMP6f average from GCaMP6f maximum projection (magenta), right panel: merge of GCaMP6f average (green) and ΔF (magenta). Bottom panels: Zoom in of boxed spine from cell in top left. First panel is GCaMP6f average, the 2^nd^ through 4^th^ panels are individual frames showing a single mSCaT at 0 msec, 400 msec, and 800 msec respectively. Red circles indicate ROIs for data traces shown in B.
B. ΔF/F traces from spine and dendrite regions circled in red in A.
C. Frequency histogram of mSCaT amplitude for individual synapses across 923 spines from 8 cells.
D. Frequency histogram of mSCaT frequency for individual synapses across 923 spines from 8 cells.
E. Representative GCaMP6f traces demonstrating that treatment with ryanodine, thapsigargin, DNQX, and nifedipine (Blockers) did not alter mSCaT amplitude compared to vehicle treatment.
F. Quantification of effect of blockers on mSCaT amplitude compared to vehicle treatment revealed that blockade of non-NMDAR sources of Ca^2+^ did not impact mSCaT amplitude.
G. Representative GCaMP6f traces demonstrating that treatment with APV eliminated mSCaTs compared to vehicle treatment.
H. Treatment with APV eliminates 94% of events.
I. Raising extracellular Ca^2+^ causes increased mSCaT amplitude, while application of 30 μM, 100 μM, and 1 mM Mg2+ reduce mSCaT amplitude. For example mSCaT traces see Extended Data Figure 1-1.

In order to confirm that our measurement is sensitive to changes in NMDAR activation, we tested the dynamic range of GCaMP6f imaging at individual synapses. We reasoned that if the indicator is not near saturation then increases in extracellular Ca^2+^ will produce approximately proportional increases in fluorescence intensity (F), otherwise very large Ca^2+^ influx will saturate the indicator and result in a disproportionately low ΔF/F. To test this, we raised the concentration of Ca^2+^ in the ACSF from 1.5 mM to 1.88 mM and measured mSCaT amplitude. We observed a significant increase in the average mSCaT amplitude (vehicle: 1.138 ± 0.031 ΔF/F, n=611/17; 1.88 mM Ca^2+^: 1.56 ± 0.051 ΔF/F, n=377/4; p<0.0001, Kruskal-Wallis) (Fig. 1I; Extended Data Figure 1-1), confirming that under our experimental conditions there was sufficient upper range to detect larger mSCaTs. To probe for sensitivity to smaller responses, and establish a lower limit to our detection, we added extracellular Mg^2+^ to block NMDAR channels, which we predicted would decrease the amplitude of mSCaTs. The IC_50_ of Mg^2+^ at a typical resting potential of −60 mV is ∼20 μM for NMDARs (Kuner and Schoepfer, 1996). Increasing the extracellular Mg^2+^ concentration from 0 to 30 μM caused a significant decrease of close to 50% in the range of mSCaT amplitudes, as expected (vehicle: 1.138 ± 0.031 ΔF/F, n=611/17; 30μM: 0.697 ± 0.046 ΔF/F, n=122/7; p<0.0001, Kruskal-Wallis) (Fig. 1I; Extended Data Figure 1-1), confirming that there is sufficient range to detect smaller mSCaTs. Additionally, the sensitivity of the events to [Mg^2+^] further confirms that mSCaTs reflect the amount of NMDAR activation. In order to evaluate the lower limit of mSCaT detection, we raised the Mg^2+^ concentration again, to 100 μM and 1 mM. Both concentrations decreased mSCaT amplitude (100 μM: 0.513 ± 0.049 ΔF/F, n=66/5; 1 mM: 0.31 ± 0.057 ΔF/F, n=20/4; p<0.0001, Kruskal-Wallis (Fig. 1I; Extended Data Figure 1-1) and decreased mSCaT frequency (Hz) (vehicle: 0.04 ± 0.002, n=414/3; 100 μM: 0.005 ± 0.0001, n=132/5; 1 mM: 0.0004 ± 0.0002, n=89/4; p<0.0001, Kruskal-Wallis). Interestingly, while the above Mg^2+^ experiments were performed in the absence of DNQX, we observed the same effects of Mg^2+^ in the presence of DNQX (data not shown). Together this suggests that under more physiological conditions, AMPAR activation after spontaneous release does not alone cause sufficient depolarization to relieve the Mg^2+^ block on the NMDARs. This decrease in mSCaT frequency suggests that either a subset of events are too small to detect when Mg^2+^ is present, or that some synaptic NMDARs are effectively entirely blocked by Mg^2+^ at these concentrations. Measuring the distribution of individual mSCaTs in elevated Mg^2+^ revealed a lower ΔF/F limit of 0.1 ΔF/F, the amplitude of the smallest events observed, below which mSCaTs cannot be reliably detected. While we cannot deduce the true amount of Ca^2+^ influx from the GCaMP6f ΔF/F due to the nonlinearities inherent in the imaging approach, these data taken together indicate a large dynamic range of the reporter, and suggest it provides an acceptable readout of the strength of NMDAR activation at individual synapses. In order to ensure that NMDARs were minimally blocked, all other experiments were performed in the absence of added Mg^2+^.

We observed that mSCaT amplitudes and frequencies were highly variable between synapses (Fig. 1C, D) with population inter-spine CVs of 0.64 and 0.99 respectively. Interestingly, the mean within-spine CV for event amplitude was also quite high, nearly at the level as what was observed between spines (0.57 ± 0.01 n=700/10, at least 3 events per spine). This variability was not due to differences in co-agonist availability, as treatment with 100 μM D-serine actually increased intra-spine mSCaT variability (mean CV normalized to baseline: vehicle: 1.05 ± 0.05, n=159/4; D-serine: 1.28 ± 0.05, n=252/4; p=0.002, unpaired t-test). We therefore asked what synaptic or receptor properties could underlie this high degree of variability in receptor activation by spontaneous release both within and between synapses.

### GluN2B-NMDARs mediate the majority of the NMDAR-dependent Ca^2+^ influx by spontaneous release

A critical regulator of NMDAR function and downstream signaling is the composition of its GluN2 subunit. Specifically, GluN2B-NMDARs are important mediators of synaptic plasticity (Foster et al., 2010; Sanhueza et al., 2011; Shipton and Paulsen, 2014a), which makes them well suited to mediate the plasticity induced by changes in spontaneous NMDAR activation. Therefore, we asked what proportion of mSCaTs in mature synapses is comprised of GluN2B- NMDARs activation. To address this, we imaged GCAMP6f before and after bath application of the GluN2B-specific antagonist ifenprodil (6 μM), or vehicle (Fig. 2A, B). Ifenprodil at this concentration is expected to nearly completely block GluN2B-NMDARs, block ∼20% of GluN2A/B-NMDARs, and have almost no effect on GluN2A-NMDARs (Stroebel et al., 2014). In our cells, treatment with ifenprodil caused a ∼35% reduction in mSCaT amplitude (normalized amplitude: vehicle: 0.977 ± 0.015, n=1204/20; ifenprodil: 0.622 ± 0.017, n=572/10; p<0.0001, Kruskal-Wallis) (Fig. 2C, D, G). Interestingly, we observed a dramatic reduction in mSCaT frequency following application of ifenprodil, with ∼50% of synapses completely silenced by ifenprodil application and an ∼81% reduction in overall mSCaT frequency (normalized frequency: vehicle: 0.82 ± 0.022, n=961/20; ifenprodil (3 wk): 0.138 ± 0.006, n=788/10; p<0.0001, Kruskal-Wallis) (Fig. 2E, F, H). The dramatic drop in event frequency along with complete blockade of events at half of the synapses suggests a significant portion of mSCaTs are entirely GluN2B-NMDAR mediated, and that GluN2A-NMDARs contribute very little to NMDAR-mediated Ca^2+^ influx in response to spontaneous release.

**Figure 2:**
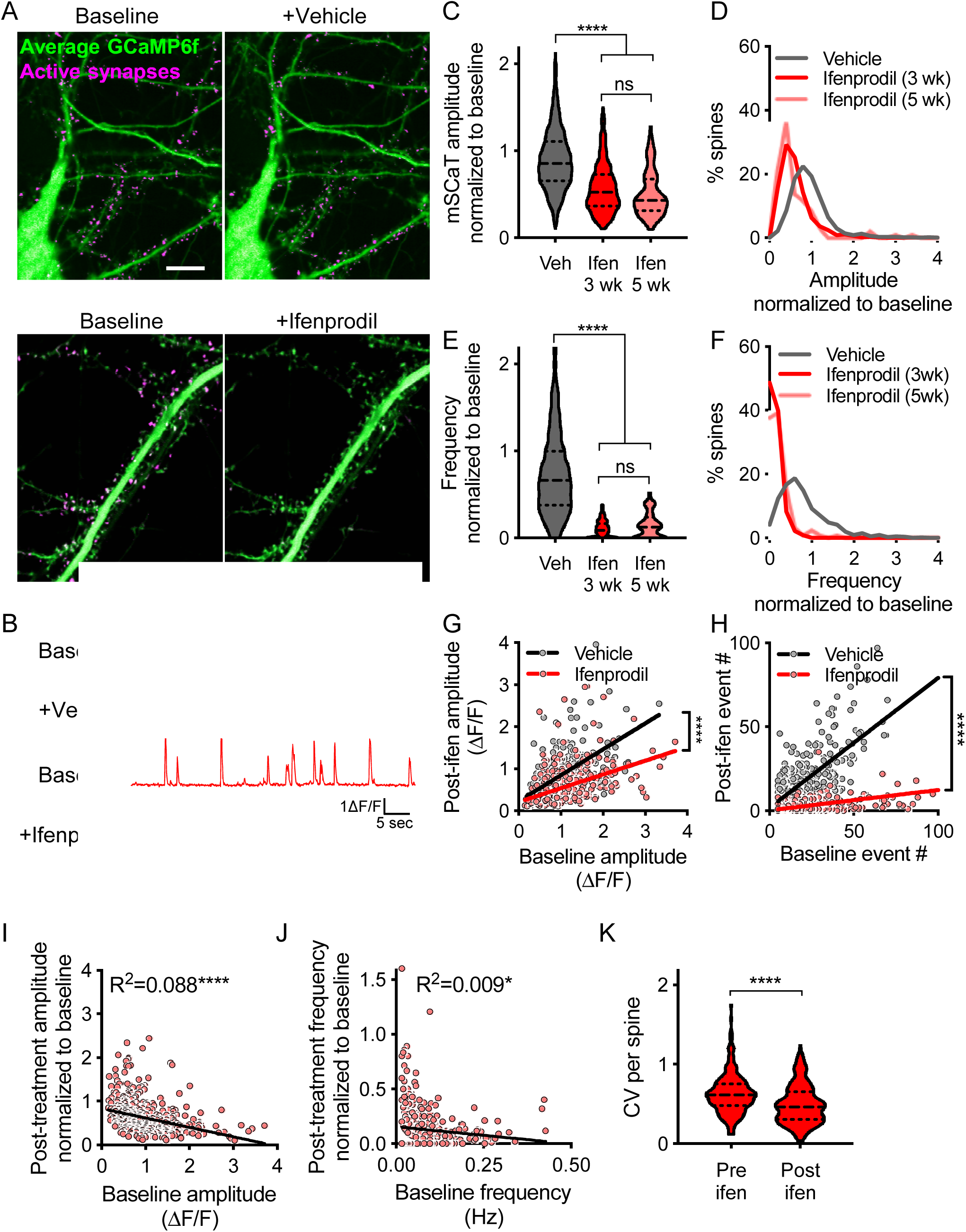
GluN2B-NMDARs mediate majority of response to spontaneous glutamate release. A. Average projection of GCaMP6f (green) at baseline and following treatment with active synapse shown in magenta. Following treatment with ifenprodil there is a clear reduction in the number of active spines compared to baseline (scale bar: 10 μm).
B. Example traces from spines treated with either vehicle (black) or ifenprodil (red).
C. Ifenprodil treatment leads to a reduction in mSCaT amplitude at both 3 and 5 weeks. Outliers removed for data display. Solid line represents median and dashed lines indicate 1^st^ and 3^rd^ quartile.
D. Cumulative probability for amplitude normalized to baseline for ifenprodil and vehicle treated cells at 3 and 5 weeks.
E. Ifenprodil treatment leads to a reduction in mSCaT frequency at both 3 and 5 weeks. Outliers removed for data display. Solid line represents median and dashed lines indicate 1^st^ and 3^rd^ quartile. See also Extended Data Figure 2-1 and 2-2.
F. Cumulative probability for frequency normalized to baseline for ifenprodil and vehicle treated cells at 3 and 5 weeks. Treatment with ifenprodil reduces mSCaT frequency with a portion of spines completely blocked.
G. Post-treatment mSCaT amplitude versus baseline mSCaT amplitude has a slope of 0.6167 ± 0.033 for vehicle treated cells and 0.327 ± 0.025 for ifenprodil treated cells and these slopes are significantly different (p<0.0001). Post-treatment mSCaT amplitude is correlated with baseline amplitude for both vehicle treated synapses (R^2^=0.42, p<0.0001) and ifenprodil treated synapses (R^2^=0.22, p<0.0001). See also Extended Data Figure 2-1 and 2-2.
H. Baseline mSCaT frequency versus post treatment mSCaT frequency reveals that nearly all synapses show a reduction in event number with ifenprodil treatment (vehicle: slope= 0.7984 ± 0.02368; ifenprodil: slope= 0.108 ± 0.005452; p<0.0001). Post-treatment mSCaT amplitude is correlated with baseline amplitude for both vehicle treated synapses (R^2^=0.56, p<0.0001) and ifenprodil treated synapses (R^2^=0.31, p<0.0001).
I. Normalized amplitude post ifenprodil treatment is negatively correlated with baseline amplitude.
J. Normalized mSCaT frequency post ifenprodil treatment is negatively correlated with baseline mSCaT frequency.
K. Within spine CV decreases following ifenprodil treatment.

Additionally, because GluN2B-NMDAR levels are developmentally regulated in some brain areas (Chen et al., 2000; Barth and Malenka, 2001; Ritter et al., 2002; Yashiro and Philpot, 2008), we imaged neurons at 5 weeks, to assess whether the large effect of ifenprodil on mSCaT frequency was due to the age of the cells. We found a significant reduction in amplitude and frequency with ifenprodil in 5-week cells (normalized amplitude (5 wk): 0.608 ± 0.049, n=94/9; normalized frequency (5 wk): 0.196 ± 0.022, n=130/9; p<0.0001, Kruskal-Wallis), and no difference between the effects of ifenprodil on 3 week or 5 week old neurons (amplitude: p= 0.546, Kruskal-Wallis; frequency: p=0.0247, Kruskal-Wallis) (Fig. 2C, D, E, F), indicating that GluN2B-NMDARs contribute significantly to spontaneous events in mature cultured hippocampal neurons of different ages. It is conceivable that expression of GCaMP6f, by chronic alteration of Ca^2+^ buffering in the cell, could lead to abnormal expression of GluN2 subunits and prevent the typical developmental shift in the ratio of GluN2B:GluN2A subunits (Yashiro and Philpot, 2008). To test this, we examined subunit expression levels at DIV 12 and 19 in control cultures, or in cultures that had been infected with GCaMP6f or GFP. In uninfected cultures, we observed the expected decrease in the ratio, and this decrease was unaltered in either virus-infected condition (Extended Data Figure 2-1C, D & Extended Data Figure 2-2). We then examined ifenprodil sensitivity in neurons that were only transiently transfected with GCaMP6f, rather than infected. We found that neurons that expressed GCaMP6f from DIV 18- 21 showed the same high ifenprodil sensitivity as those chronically expressing via virus infection (Normalized amplitude: Veh-infected: 0.98 ± 0.016, n=1204/20; Ifen-Infected: 0.622 ± 0.017, n=572/10; Veh-transfected: 1.07 ± 0.035, n=103/4; Ifen-transfected: 0.633 ± 0.051, n=81/4; p<0.0001, one-way ANOVA; Normalized frequency: Veh-infected: 0.82 ± 0.022, n=961/20; Ifen-Infected: 0.137 ± 0.006, n=788/10; Veh-transfected: 0.855 ± 0.063, n=110/4; Ifen-transfected: ± 0.048, n=142/4; p<0.0001, one-way ANOVA) (Extended Data Figure 2-1A,B). These observations suggest that chronic GCaMP6f expression does not alter subunit expression.

Returning to further analysis of virally infected neurons, we also observed that synapses with larger mSCaTs at baseline were associated with a greater effect of ifenprodil on mSCaT amplitude (R^2^= 0.089, p<0.0001, n=544/10) (Fig. 2I), and spines with higher mSCaT frequency at baseline tended to have a larger portion of events blocked with ifenprodil (R^2^= 0.012, p=0.003, n=751/10) (Fig. 2J). Thus, more active synapses, with more NMDAR activation, have a larger contribution of GluN2B-containing NMDARs to their events. Because GluN2B-NMDARs have longer open times and lower open probability than GluN2A-NMDARs, it is likely that there is increased variability in Ca^2+^ influx through GluN2B-NMDARs (Santucci and Raghavachari, 2008). Indeed, we observed that mSCaT amplitude variance was slightly reduced following ifenprodil treatment (baseline CV: 0.633 ± 0.02; post-ifenprodil CV: 0.495 ± 0.016; n=225, p<0.0001, paired t-test, Fig. 2K), however this could at least be partially due to the large reduction in the number of events. Overall, NMDAR subtype may contribute to the high degree of variability in NMDAR-mediated Ca^2+^ influx. Taken together, these results not only indicate that GluN2B-NMDARs contribute to events at these synapses, but also that a significant portion of mSCaTs are largely GluN2B-NMDAR mediated.

### NMDAR activation is independent of spine and synapse size

While GluN2B-NMDARs contribute to variability of mSCaT amplitude within synapses, other parameters may mediate variability between synapses. While there is evidence that the number of NMDARs present in the PSD (Kharazia et al., 1999; Takumi et al., 1999; Shinohara et al., 2008; Chen et al., 2015) as well as the magnitude of their activation by evoked release is not related to spine size (Nimchinsky et al., 2004), there is a strong relationship between spine size and the amount of activation of other receptor types, particularly AMPARs (El-Husseini et al., 2000; Masanori Matsuzaki et al., 2004; Araki et al., 2015). Since NMDAR activation by spontaneous release may be regulated distinctly from NMDAR activation by evoked release, we tested the relationship between mSCaTs and spine size. We measured spine area at individual synapses based on post-Ca^2+^ imaging z-stacks (Fig. 3A, B). Spine area ranged from 0.163 μm^2^ to 1.411 μm^2^ with a mean spine area of 0.639 ± 0.009 μm^2^. By matching spine area to mSCaT data for each synapse, we found that mean mSCaT amplitude per spine was weakly negatively correlated with spine area for spontaneous release events (R^2^=0.008, p=0.028, n=628/10). A similar result was seen with binned spine area data to reduce overpowering the analysis (R^2^=0.256, p=0.023, n=20/10 (Fig. 3C). However, this negative correlation is likely due to smaller proportional Ca^2+^ influx in larger spines rather than differences in the amount of NMDAR activation (Nimchinsky et al., 2004). Overall, this suggests that NMDAR activation does not substantially scale with spine size.

**Figure 3:**
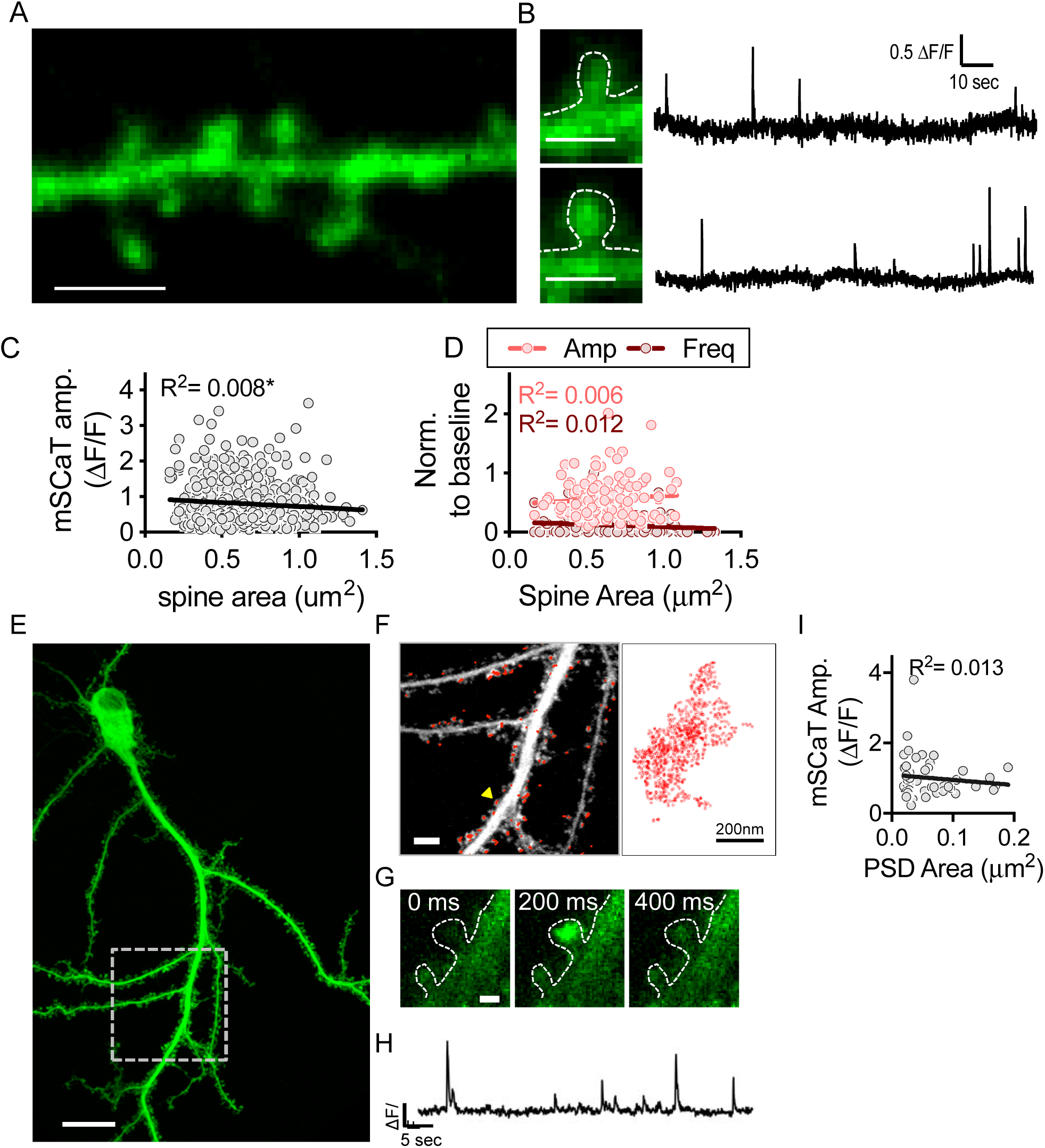
NMDAR activation is independent of spine area. A. Example stretch of dendrite from post-Ca^2+^ imaging GCaMP6f z-stack (scale bar 10 μm).
B. Zoom in of spines indicated by white arrowheads from A paired with their respective Ca^2+^ traces (scale bar 1 μm).
C. mSCaT amplitude weakly, negatively correlates with spine area. First plot is all spines, second plot binned data.
D. Effect of ifenprodil on amplitude (light red) and frequency (dark red) does not correlate with spine area. Left plot is all spines, right plot is binned data.
E. GCaMP6f max projection acquired directly following Ca^2+^ imaging (scale bar is 20 μm) of dSTORM imaged neuron. White box indicates area where super-resolution imaging was performed.
F. Zoom in on the region from E. Max projection of GCaMP6f stack acquired at time of STORM imaging (white) (scale bar is 5 μm). Super-resolved shank localizations are shown in red. Right is a zoom-in of the shank localizations from the spine indicated with the yellow arrowhead.
G. Zoom in on a mSCaT in the spine indicated by yellow arrowhead in F.
H. Ca^2+^ trace from spine indicated by yellow arrowhead in F.
I. Amplitude does not correlate with PSD area.

GluN2B-NMDARs have been shown to exit synapses that have undergone LTP (Dupuis et al., 2014) and those synapses tend to be larger (Matsuzaki et al., 2001; Malenka and Bear, 2004; Matsuzaki et al., 2004). Therefore, it is possible there is a smaller contribution of GluN2B- NMDARs at larger spines. To address this, we examined the correlation between the effect of ifenprodil on mSCaTs at individual synapses and spine area. We found that spine area did not correlate with the magnitude of ifenprodil blockade on amplitude or frequency of mSCaTs (normalized amplitude: raw: R^2^=0.006, p=0.388, n=127/6; binned: R^2^=0.050, p=0.342, n=20/6; normalized frequency: R^2^=0.012, p=0.109, n=221/6; binned: R^2^=0.059, p=0.298, n=20/6) (Fig. 3D). These data suggest that the contribution of GluN2B-NMDARs per synapse is not related to spine area.

Even though spine size is often related to synapse size (Harris et al., 2014), the PSD is much smaller than the spine itself, and is a highly dynamic structure that is strongly correlated to the size of the active zone across the cleft (Harris and Stevens, 1989; Schikorski and Stevens, 1997; Inoue and Okabe, 2003; MacGillavry et al., 2013). Thus, to measure the area of the PSD itself using super-resolution microscopy, we turned to dSTORM imaging of the postsynaptic protein Shank. We matched mSCaT data to super-resolved PSDs for 47 synapses from 6 neurons that were subjected to live GCaMP6f imaging followed by fixation and anti-Shank immunocytochemistry and dSTORM imaging (Fig. 3E, F, G, H). The mean PSD area was 0.060 μm^2^ ± 0.006, which is near the mean reported by electron microscopy of hippocampal synapses (mean ∼0.069 μm^2^) (Harris and Stevens, 1989; Schikorski and Stevens, 1997; Shinohara et al., 2008). We found that mSCaT amplitude was not correlated with PSD area (R^2^=0.013, p=0.452, n= 47/6) (Fig. 3I).

Taken together, this set of observations indicates that the size of the activated pool of NMDARs at individual synapses is independent of synapse size and that variability in the magnitude of NMDAR activation observed between synapses for spontaneous release events is controlled by factors other than the size of the synapse itself.

### mSCaT amplitude is correlated with synapse distance from the soma

Another potential source of variability in mSCaT amplitude and frequency is the position of the synapse within the dendritic tree. In fact, NMDAR-mediated Ca^2+^ influx has been shown to be increased in synapses further from the cell body (Walker et al., 2017). We asked whether NMDAR activation by spontaneous glutamate release varies throughout the dendritic arbor as a function of distance from the soma or the number of branch points away from the soma (branch depth). We mapped neuronal morphology and identified spines along dendrites from z-stacks of GCaMP6f expressing neurons using semi-automatic neuron tracing and spine detection in Imaris (Fig. 4A, B, C) and identified spines in the traced image that had been Ca^2+^ imaged (Extended Data Figure 4-1). Mean mSCaT amplitude per spine was significantly correlated with distance from the soma (R^2^=0.014, p=0.036 n=316/9) (Fig. 4C), as expected (Walker et al., 2017). Interestingly, even though branch depth was correlated with distance from the soma (R^2^=0.022, p<0.0001; data not shown), branch depth did not correlate with mean mSCaT amplitude (raw data: R^2^= 0.0007, p=0.649, n=287/9) (Fig. 4E). In order to compare spines that are farther than ∼200 μm away from the soma, we repeated these experiments using a lower power objective to increase the field of view. This allowed for a larger field of view and measurement of distances up to ∼450 μm away from the soma. We observed that even at these longer distances, there was a correlation between mSCaT amplitude and distance from soma (R^2^=0.041, p<0.0001 n=746/6) (Fig. 4D, E). Interestingly, when mSCaT amplitude was compared to branch depth at longer distances there was a correlation between mSCaT amplitude and branch depth (R^2^=0.018, p=0.0002 n=746/6) (Fig. 4H,I). These results demonstrate the amount of NMDAR activation following spontaneous single vesicle release is related to the synapse distance from soma, and branch depth at more distal branches.

**Figure 4:**
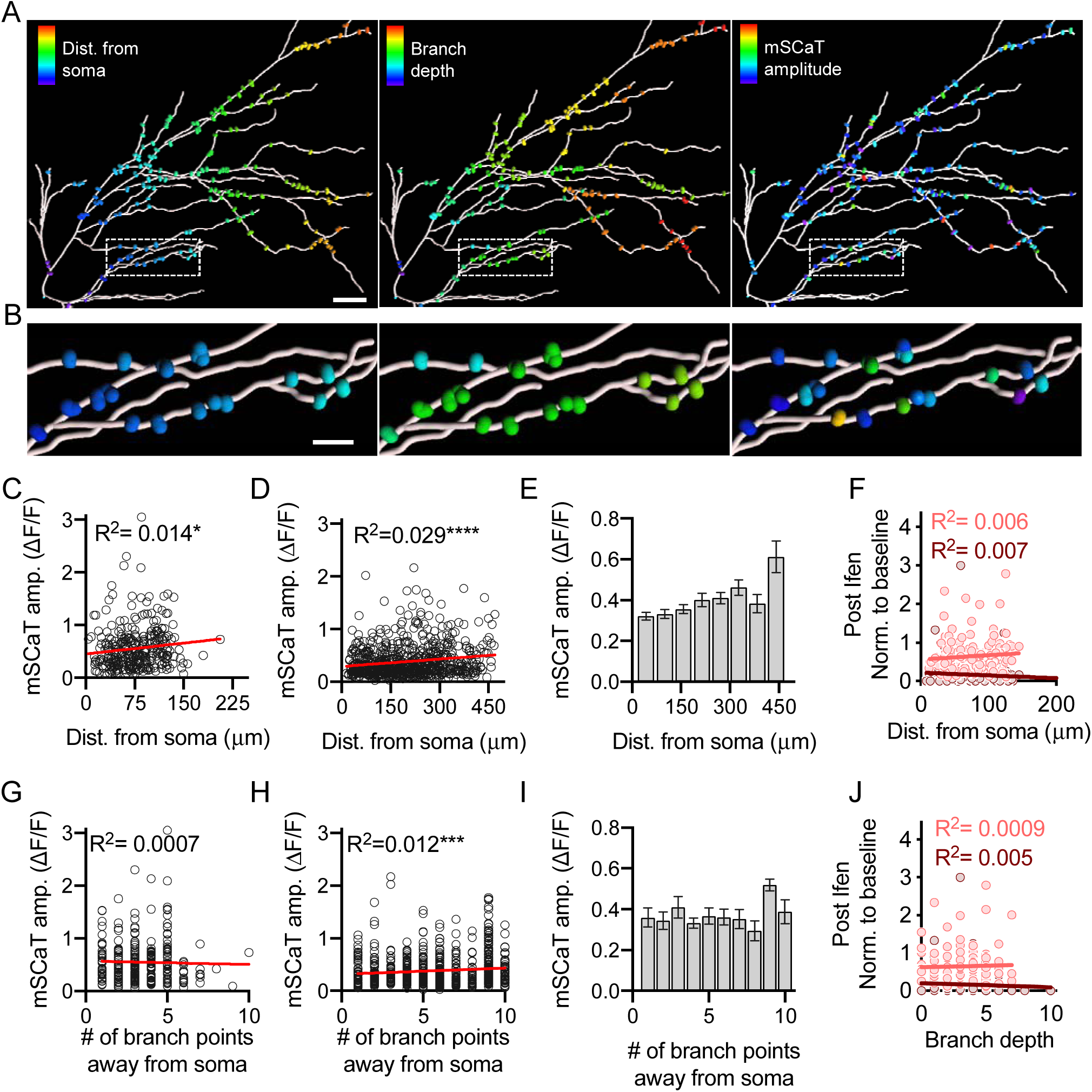
Mapping synapse position and mSCaT characteristics with Imaris. A. Ca^2+^ imaged spines along traced dendrite from Imaris. Scale bar is 30 μm. First panel shows distance from soma, second panel shows branch depth, and third panel shows mSCaT amplitude. Colors are warmer as spines that are farther from the cell body, have higher branch depth, or have larger mSCaT mean amplitudes respectively. Further details on tracing and spine identification can be seen in Extended Data Figure 4-1.
B. Zoom-in of boxed areas from A.
C. mSCaT amplitude did correlate with distance from soma for proximal spines.
D. mSCaT amplitude did correlate with distance from soma when distal spines are included.
E. Binned data from D demonstrating relationship between spine distance from the soma and mSCaT amplitude
F. Magnitude of ifenprodil effect on mSCaT amplitude (light red) and frequency (dark red) does not correlate with distance from the soma.
G. mSCaT amplitude did not correlate with the number of branch points away from the soma the synapse is (branch depth) for proximal spines.
H. mSCaT amplitude does correlate with branch depth when distal spines are included.
I. Mean mSCaT amplitude at each branch depth.
J. Magnitude of ifenprodil effect on mSCaT amplitude (light red) and frequency (dark red) did not correlate with branch depth.

We then further asked whether GluN2B-NMDAR content varied based on synapse position. We found no correlation between the magnitude of the effect of ifenprodil with either distance from the soma (normalized frequency: raw data: R^2^=0.007, p=0.237, n=209/9; binned data: R^2^=0.036, p=0.422, n=20/9; normalized amplitude: raw data: R^2^=0.006, p=0.367, n=132/9; binned data: R^2^=0.034, p=0.434, n=20/9) (Fig. 4F) or branch depth (normalized frequency: raw data: R^2^=0.005, p=0.315, n=209/9; binned data: R^2^=0.007, p=0.721, n=209/9; normalized amplitude: raw data: R^2^=0.0009, p=0.729, n=20/9; binned data: R^2^=0.00004, p=0.978, n=20/9) (Fig. 4J). Therefore, the amount of GluN2B-NMDAR activation is independent of synapse position within the dendritic arbor.

Overall, synapse position did loosely correlate with mSCaT amplitude, whereas spine and synapse size did not. Furthermore, the GluN2B component of mSCaTs is also independent of synapse size and position. Therefore, it likely that there are other factors besides size or position is responsible for the dramatic variability in spontaneous NMDAR activation between synapses.

## Discussion

In the present study, we used an all-optical approach to characterize NMDAR activation following AP-independent (spontaneous) vesicle exocytosis at individual synapses of cultured hippocampal neurons. While GCaMP6f has frequently been used to measure mSCaTs (Andreae and Burrone, 2015; Sinnen et al., 2016; Tang et al., 2016; Walker et al., 2017), we clarified that it offers a large dynamic range and permits detailed analysis of the magnitude of receptor activation. Using this approach, we found that at nearly all synapses in this preparation, GluN2B-NMDARs are the major NMDAR subtype activated during spontaneous synaptic transmission. Additionally, we observed a surprising degree of variability in mSCaT amplitude both between and within synapses. In fact, the variation at single spines was comparable to the between-spine variation, which suggests that Ca^2+^ influx through the receptor may be surprisingly independent of synapse-specific features such as the number of receptors present at the synapse. Indeed, we demonstrated that spine size, PSD area, and synapse position have a relatively small impact on mSCaTs. Therefore, this high degree of variability is likely to be dominated by differences in release position, variations in the amount of glutamate per vesicle, and random fluctuations in channel open time.

We found that even in the absence of an AMPAR antagonist, application of 1 mM Mg^2+^, which is still below the physiological concentration, nearly eliminates all observable mSCaTs. This suggests that under more physiological conditions, in the absence of other activity, many spontaneous release events do not lead to sufficient membrane depolarization to relieve the Mg^2+^ block on NMDARs, and thus result in little or no Ca^2+^ influx. Recent evidence demonstrates that glutamate binding to NMDARs can induce conformational changes in the receptor which lead to metabotropic signaling even in the absence of ion flux (Nabavi et al., 2013; Dore et al., 2016; Dore et al., 2017). Importantly, this type of NMDAR activation can mediate some forms of plasticity (Kessels et al., 2013; Aow et al., 2015; Dore et al., 2015; Stein et al., 2015; Wong and Gray, 2018). Thus, our data suggest that under physiological conditions, effects of NMDAR activation by single-vesicle release events could be mediated principally by non-ionotropic functions rather than via Ca^2+^ influx.

Our data demonstrate that GluN2B-NMDARs contribute significantly to spontaneous events at all synapses and, at roughly half of synapses, are the primary NMDAR type activated by spontaneous release. This was somewhat surprising given the prevailing notion that GluN2B- NMDARs are not present at synapses after early development (Chen et al., 2000; Barth and Malenka, 2001; Ritter et al., 2002; Yashiro and Philpot, 2008). However, this is consistent with other reports (Sinnen et al., 2016; Walker et al., 2017) and a large and growing amount of evidence suggests that GluN2B-NMDARs are found at mature hippocampal synapses (Kellermayer et al., 2018) and they contribute significantly to synaptic events (Gray et al., 2011; Xiao et al., 2016; Levy et al., 2018). Thus, while it is clear that in some brain areas a developmental switch in NMDAR subtype is pronounced, in the hippocampus it is not as prominent. Spontaneous NMDAR activation has specialized functions within the synapse (Sutton et al., 2004; Sutton et al., 2006; Kavalali et al., 2011; Andreae and Burrone, 2015), and it is possible that these functions are specifically driven by GluN2B-NMDAR activation, rather than NMDAR activation in general. It will be important to assess whether GluN2B-NMDARs are required for the downstream signaling induced by spontaneous release.

We observed not only a large contribution of GluN2B-NMDARs to mSCaTs, but also a striking lack of Ca^2+^ influx mediated by GluN2A-NMDARs. Data regarding the contribution of NMDAR subtype to synaptic responses to spontaneous glutamate release has been mixed. In previous reports, knockout of GluN2A-NMDARs reduced or eliminated NMDA-mEPSCs in mature midbrain synapses (Townsend et al., 2003; Zhao and Constantine-Paton, 2007), thus suggesting the GluN2A-NMDARs are the principle responders to spontaneous release. This may be a region-specific effect, or, since GluN2A-NMDARs are essential for normal development of synapses in many brain regions (Gambrill and Barria, 2011; Gray et al., 2011; Kannangara et al., 2014), it is possible that global GluN2A knockout alters synapses in other, unexpected ways.

While GluN2A-NMDARs did not contribute significantly to mSCaTs here, they are present at hippocampal synapses in culture and function in evoked neurotransmission (MacGillavry et al., 2013; Xiao et al., 2016; Kellermayer et al., 2018). Why they are less activated by spontaneous release? One option is that there is segregation of NMDARs such that the receptors activated by spontaneous release form a distinct pool, either a subset of or separate from those activated by evoked release. There is evidence to support this idea (Atasoy et al., 2008; Reese and Kavalali, 2016), and it possible that NMDAR subtype is specific to one pool or the other. A potential mechanism for restricting NMDAR activation is spatial segregation within the synapse. Indeed, within synapses, receptors are found to have a distinct nanoscale organization with subsynaptic high-density nanoclusters of postsynaptic proteins and AMPARs as well as GluN2B-NMDARs (Fukata et al., 2013; MacGillavry et al., 2013; Nair et al., 2013). And importantly, these postsynaptic nanodomains are aligned with presynaptic evoked release sites (Tang et al., 2016). Recently it has been demonstrated that within individual synapses, GluN2A- and GluN2B-NMDARs form distinct nanodomains that differ in size, number, and internal density (Kellermayer et al., 2018), further consistent with the idea that nano-organization could influence which receptor type is activated. Interestingly because of their biophysical properties, GluN2B-NMDARs are especially likely to be sensitive to their positioning with respect to the site of release; receptors within ∼50 nm of the site of release three times more likely to open than those located ∼200 nm from the site of release (Santucci and Raghavachari, 2008). Therefore, the position of release site with respect to NMDARs may restrict not just the total amount of NMDAR activation but also which synaptic NMDARs are able to be activated by spontaneous release. Further, mapping of release sites during spontaneous and evoked release revealed that these release modes display different spatial patterns within the active zone (Tang et al., 2016). Given the possibility that GluN2B-NMDARs are positioned within the synapses as to be relevant for spontaneous release, the contribution of GluN2B- NMDARs to events in mature synapses may have been underestimated due to a focus on evoked release.

In addition to GluN2B-NMDARs and GluN2A-NMDARs, there is also a significant population of triheteromeric receptors (GluN2A/B-NMDARs) thought to be at mature hippocampal synapses (Rauner and Köhr, 2011; Paoletti et al., 2013; Tovar et al., 2013; Stroebel et al., 2018). Unfortunately, these are difficult to study in situ due to a lack of specific pharmacological agents. Based on dose-inhibition curves for ifenprodil for the different receptor subtypes, the ifenprodil concentration utilized here blocked nearly all GluN2B-NMDAR activation (IC_50_: 0.15 µM), but also ∼20% of any GluN2A/B-NMDAR-mediated response, though is expected to have had no impact on GluN2A-NMDARs (IC_50_: >20 µM) (Paoletti, 2011; Hansen et al., 2014; Stroebel et al., 2014), Though it is difficult with existing reagents to specify how prominent their role is, triheteromeric receptors thus probably contribute to some spontaneous events. However, based on the dramatic reduction in the number of events with ifenprodil treatment, it is likely that GluN2B-NMDARs mediates the majority of the Ca^2+^ influx due to spontaneous release.

We observed a high degree of variability in mSCaT amplitude both between and within synapses. Differences in the number of receptors activated per event or the NMDAR subtype activated could underlie variability in event amplitude. If the number of NMDARs activated per event was dominating this inter-event variability, then this would lead to the prediction that some synapses with more NMDARs, or a higher density of NMDARs apposed to release sites would have overall larger events. One consequence of this would be that the variability within a single synapse would be smaller than the variability between synapses. However, we observed a similar amount of mSCaT amplitude variability within synapses as between synapses, thus difference in the number of NMDARs able to be activated between synapses is not likely the dominant source of variability. GluN2B-NMDARs have longer open times and longer burst duration than GluN2A-NMDARs but have a much lower open probability, suggesting that they may contribute more to variability in the amount of Ca^2+^ influx per event (Santucci and Raghavachari, 2008). In fact, in single-channel recordings, ifenprodil reduces variability of NMDAR total open time (Pina-Crespo and Gibb, 2002). Consistent with this prediction, we observed there was a decrease in mean CV of mSCaT amplitude from 0.6 to 0.5 following ifenprodil treatment. This suggests that variability in Ca^2+^ influx through GluN2B-NMDARs substantially contributes to the differences in mSCaT amplitude between events, and that Ca^2+^ influx through activated GluN2A-NMDARs is less variable.

Spine size was not substantially correlated with the amount of Ca^2+^ influx per event. Similarly, NMDAR activation does not scale with spine size following glutamate uncaging (Sobczyk et al., 2005; Takasaki and Sabatini, 2014) or evoked release (Nimchinsky et al., 2004). Together, these observations indicate that the amount of NMDAR activation is independent from spine size for both release modes. Another spine feature that may alter mSCaT amplitude and contribute to variability is spine neck diameter, since this could alter Ca^2+^ retention in the spine (Svoboda et al., 1996) Additionally, while we did not measure a strong correlation between spine size and mSCaT amplitude, it is nevertheless possible that small fluctuations in spine area during the course of the experiment could contribute slightly to the variability observed in mSCaT amplitude. Furthermore, PSD size measured with correlative super-resolution imaging was also unrelated to NMDAR activation. We did observe a weak negative correlation between spine size and mSCaT amplitude, however, we suspect that this is due to a decrease not in actual Ca^2+^ influx, but in the ratio of Ca^2+^ influx to total GCaMP6f in the compartment (that is, very large spines have a large basal F).

It is worth highlighting this novel combination of super-resolution imaging with functional measures at individual synapses. Nanoscale protein organization is hypothesized to control many aspects of synaptic function and signaling (Bourne and Harris, 2012; Choquet and Triller, 2013; Biederer et al., 2017; Chamma and Thoumine, 2018), so this correlative approach will be important for future tests of how NMDAR activation is impacted by other nanoscale synaptic features, such as the presence of subsynaptic scaffold nanoclusters (Fukata et al., 2013; MacGillavry et al., 2013; Nair et al., 2013; Broadhead et al., 2016; Tang et al., 2016). Rapidly improving and diversifying sensors for neurotransmitters and intracellular messengers (Marvin et al., 2013; Mehta et al., 2018; Ross et al., 2018), also suggest that it will be possible to more directly measure the link between synapse nanostructure and both the ionotropic and non-ionotropic activity of the receptor. Conversely, synaptic function can alter subsynaptic protein distributions, potentially mediating aspects of functional plasticity (Glebov et al., 2017; Chen et al., 2018). We anticipate the correlative approach will be integral for exploration of dynamic synaptic signaling.

In another series of experiments, we asked whether synapse distance from the soma or degree of branch complexity plays a role in variability in the amount of NMDAR activation. Previous work has suggested that NMDAR-mediated Ca^2+^ influx is larger at synapses further away from the soma (Walker et al., 2017). Our data was consistent with this finding and revealed a significant correlation of mSCaT amplitude with the distance of the synapse from the soma, but not with branch complexity. However, there is evidence that spine size is correlated with distance from the soma and that more distal synapses tend to be smaller than those more proximal (Katz et al., 2009; Walker et al., 2017). Therefore, the relationship between spine distance and event amplitude may in part arise simply from a larger proportional Ca^2+^ influx at a subset of distal spines that are much smaller than proximal spines. Additionally, we found that the amount of blockade with ifenprodil was unrelated to synapse position. Overall, we conclude that despite a slight correlation between synapse position and mSCaT amplitude, synapse position is not a primary modulator of NMDAR-mediated Ca^2+^ influx between synapses.

Major remaining options for the source of variability in NMDAR activation are phosphorylation state of NMDARs, presynaptic changes in the amount of glutamate released between events, or finally, random variation in channel open time. Phosphorylation of the NMDAR can modify channel conductance and lead to changes in Ca^2+^ influx (Wang and Salter, 1994; Salter and Kalia, 2004; Skeberdis et al., 2006; Chen and Roche, 2007). These changes can occur on the timescale of minutes (Wang and Salter, 1994), which while not completely incompatible with our average mSCaT frequency of ∼2 events per minute, is unlikely to have contributed substantially to within-synapse variance in our experiments. However, it has been shown that preventing receptor phosphorylation alters basal transmission (Wang and Salter, 1994; Skeberdis et al., 2006), which suggests that there is some basal level of NMDAR phosphorylation ongoing that can modify NMDAR properties. Thus, while it is unclear whether NMDAR receptor phosphorylation would be sufficient to mediate the large amount of variability observed within synapses, it appears able to play a role in distinguishing NMDAR-mediated Ca^2+^ influx between synapses.

Differences in the amount of glutamate release per event have long been thought to be a major contributor to variability in the amount of receptor activation by spontaneous release (Bekkers et al., 1990; Liu et al., 1999; McAllister and Stevens, 2000; Hanse and Gustafsson, 2001; Franks et al., 2003). Modeling the amount of AMPAR activation with varying quantal size has demonstrated an increase in the number of activated receptors as glutamate molecules per vesicle is increased (Franks et al., 2003). Despite the likely small number of NMDARs activated per event (Nimchinsky et al., 2004), it is possible that there is variability in the number of NMDARs activated that is in part due to differences in the amount of glutamate released at each event. In addition to the amount of glutamate per vesicle, the release site position with respect to receptor position could also underlie variability in the amount of receptor activation. Modeling receptor activation due to glutamate release at different distances from the highest density of receptors has suggested that release site position could play a large role in the variability in response amplitude (Uteshev and Pennefather, 1996; MacGillavry et al., 2013; Savtchenko and Rusakov, 2013). Especially in the case of spontaneous release which may occur more randomly across the active zone than evoked release (Tang et al., 2016), release position may play a role in inter-event variability. Finally, another likely source of variability in Ca^2+^ influx per event is stochastic channel open-close transitions (Franks et al., 2003). GluN2B-NMDARs have a relatively low open probability (Chen et al., 1999), which could produce large essentially random changes in the amount of Ca^2+^ influx per event (Yeung et al., 2004; Zeng and Holmes, 2010). Especially in this case where there are very few NMDARs activated per event, these random fluctuations could dominate the variability.

Overall, we conclude that the high degree of variability in the amount of Ca^2+^ influx through spontaneous activated NMDARs is not primarily due to synapse-specific features including the number of available receptors, NMDAR subtype, synapse size, or synapse position within the dendritic tree. Rather, the high degree of variability of spontaneous NMDAR activation is most likely dominated by nanoscale intrinsic properties of the synapse that influence receptor activation probability, including receptor position, release site position, and the glutamate concentration profile following each release event, and by the highly varying open time of the channels that do become activated.

## Acknowledgements

We thank Sai Sachin Divakaruni, Aaron Levy, and other members of the Blanpied Lab for helpful discussion, Minerva Contreras for invaluable technical assistance and the University of Maryland School of Medicine Center for Innovative Biomedical Resources Confocal Microscopy Facility.

---

**Figure 1-1:**
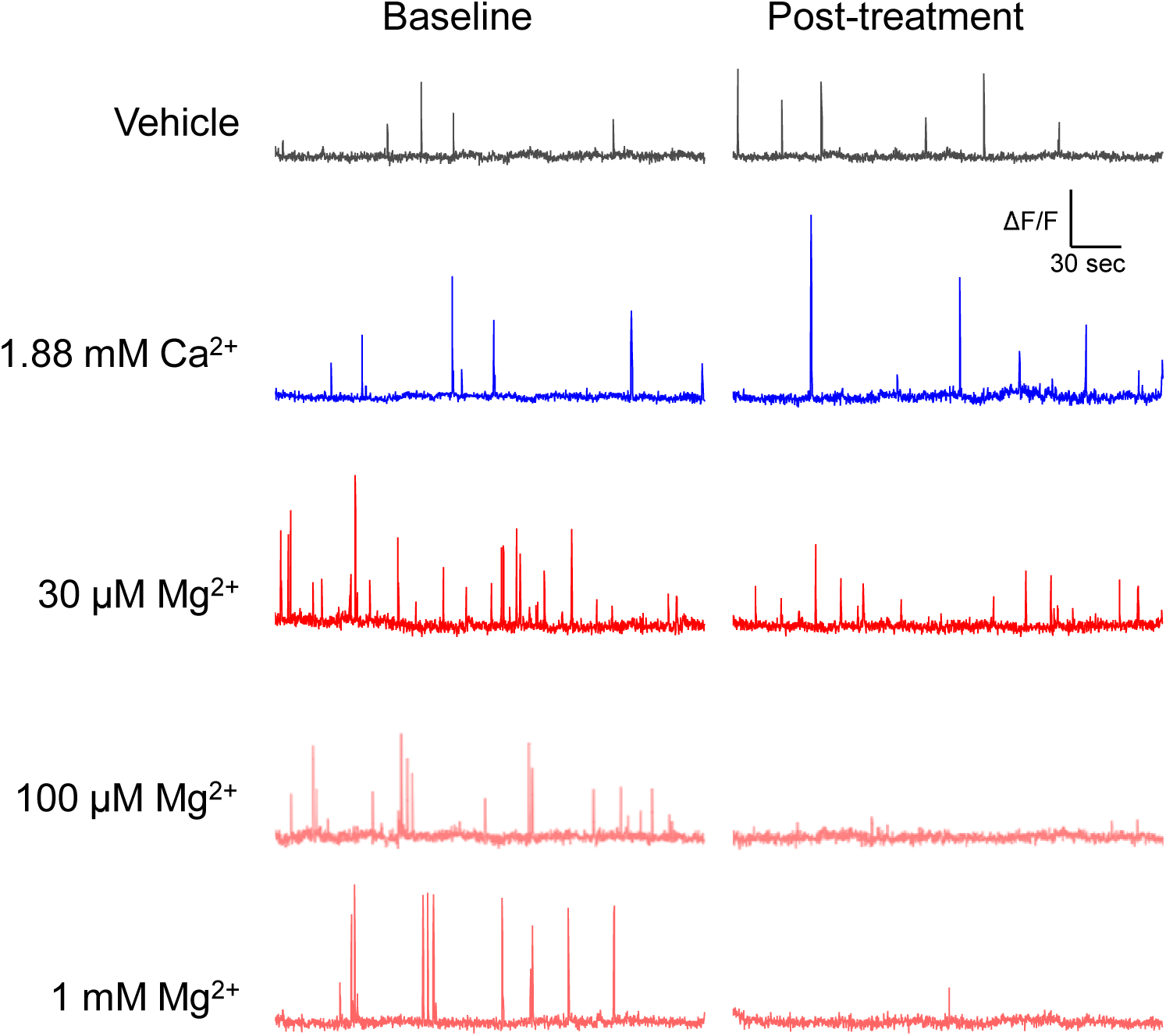
Example mSCaT traces from cells in Figure 1. Example mSCaT traces for data from Figure 1I.

**Figure 2-1:**
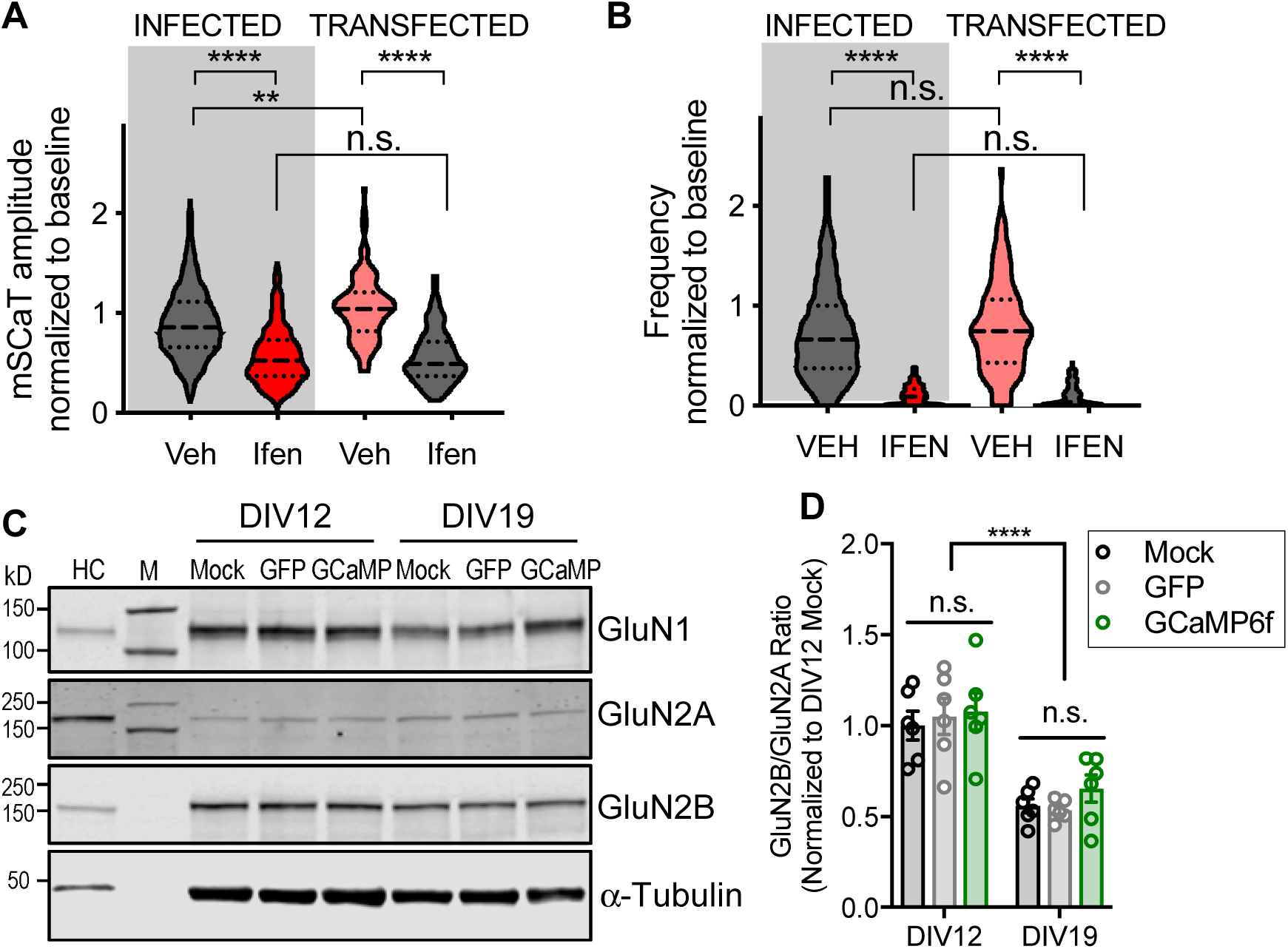
Chronic expression of GCaMP6f does not alter GluN2B-NMDAR contribution to mSCaTs or GluN2B developmental shift. A. Ifenprodil has the same effect on mSCaT amplitude for transiently transfected cells as infected cells (Normalized amplitude: Veh-infected: 0.98 ± 0.016, n=1196/20; Ifen-Infected: 0.622 ± 0.017, n=565/10; Veh-transfected: 1.07 ± 0.035, n=103/4; Ifen-transfected: 0.633 ± 0.051, n=81/4; p<0.0001, one-way ANOVA).
B. Ifenprodil has the same effect on mSCaT frequency for transiently transfected cells as infected cells (Normalized frequency: Veh-infected: 0.82 ± 0.022, n=961/20; Ifen-Infected: 0.137 ± 0.006, n=788/10; Veh-transfected: 0.855 ± 0.063, n=110/4; Ifen-transfected: 0.20 ± 0.048, n=142/4; p<0.0001, one-way ANOVA).
C. Hippocampal neurons were infected with AAV-GFP or AAV-GCaMP6f on DIV0 and harvested on DIV12 or DIV19. Representative immunoblots for GluN1, GluN2A, GluN2B, and -Tubulin are shown. Molecular weight markers (kD) indicated for each immunoblot. Sample from adult rat hippocampus (HC) included as positive control.
D. Quantitation shows decrease in GluN2B/GluN2A ratio from DIV12 to DIV19. Graph depicts normalized mean GluN2B/GluN2A ratios ± SEM (DIV factor: p<0.0001; Infection factor p=0.5009; Interaction: p=0.812; two-way ANOVA).

**Figure 2-2:**
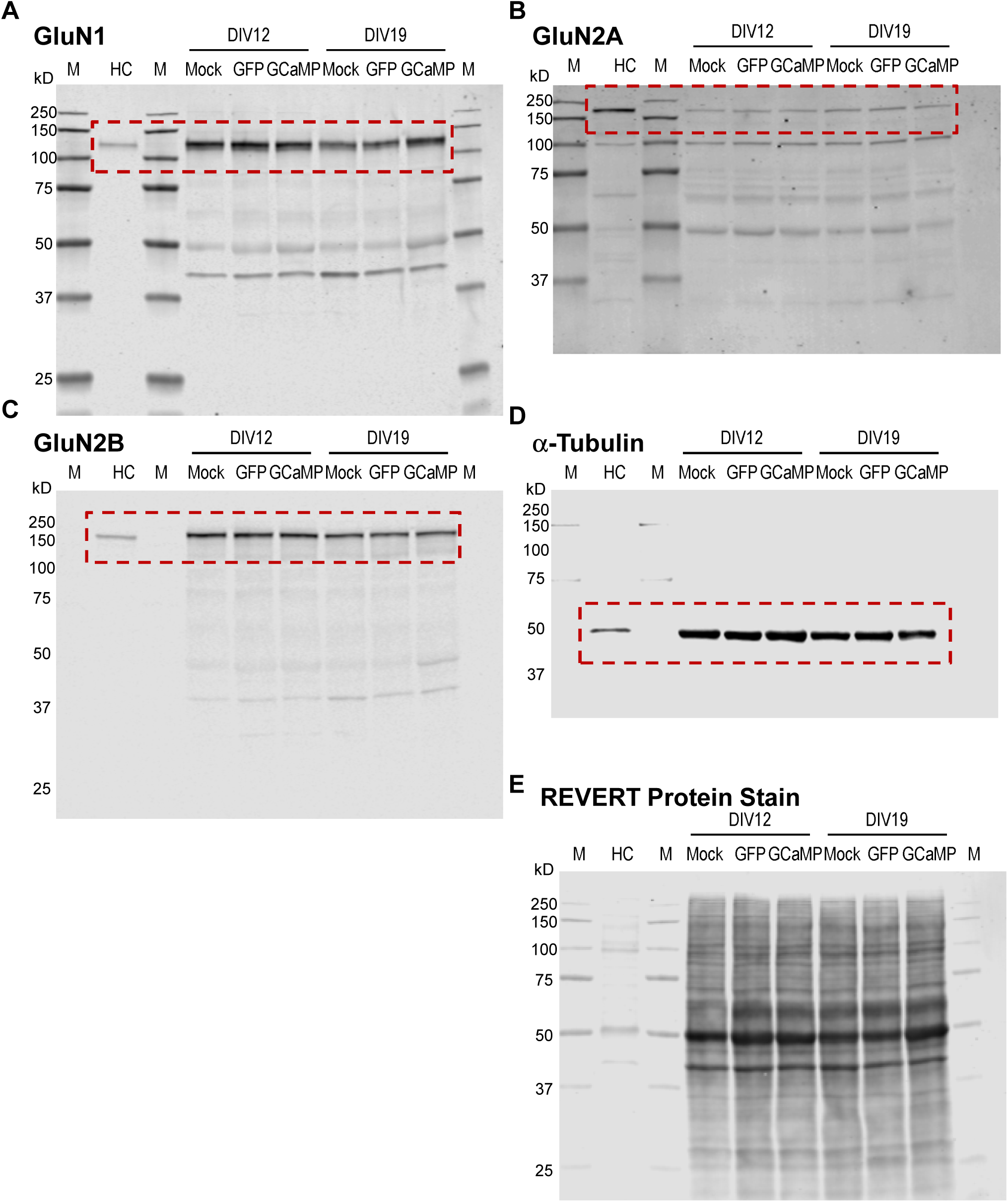
Representative full immunoblots from Figure 2-1. A.-D. Representative full immunoblots from Figure 2-1.
E. Full blot of REVERT protein stain.

**Figure 4-1:**
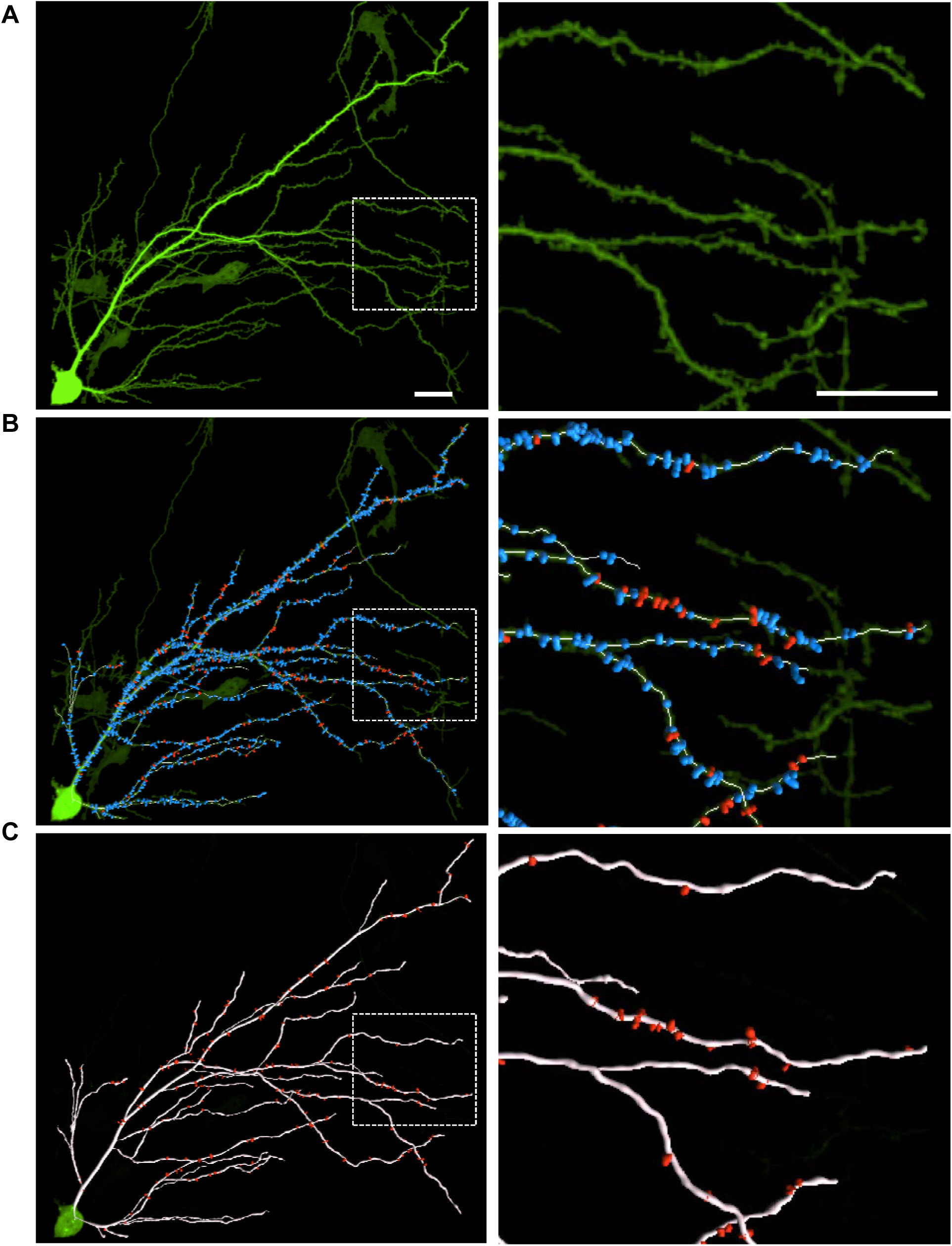
Imaris tracing examples. A. Max projection of GCaMP6f stack taken following GCaMP6f imaging. Zoom in of white box is on the right. Scale bars are 30 μm.
B. Semi-automatic spine detection on max projection of GCaMP6f. Blue spines are spines that do not have Ca^2+^ imaging data while spines in red do. Zoom in of white box is on the right. Scale bars are 30 μm.
C. Semi-automatic dendrite and spine detection in Imaris detects dimensions of cell features based on fluorescence.

